# Analysis of biological networks using Krylov subspace trajectories

**DOI:** 10.64898/2026.03.29.715092

**Authors:** H. Robert Frost

## Abstract

We describe an approach for analyzing biological networks using rows of the Krylov subspace of the adjacency matrix. Specifically, we explore the scenario where the Krylov subspace matrix is computed via power iteration using a non-random and potentially non-uniform initial vector that captures a specific biological state or perturbation. In this case, the rows the Krylov subspace matrix (i.e., Krylov trajectories) carry important functional information about the network nodes in the biological context represented by the initial vector. We demonstrate the utility of this approach for community detection and perturbation analysis using the C. Elegans neural network.

## 1 Introduction

Network-based methods are widely used for the analysis of biological data [1]. In some cases, the biological system has a physical network structure (e.g., axon connections between neurons in the brain [2]) but in many cases the network model is an abstraction of pair-wise, and potentially directed, relationships between biological entities (e.g., gene regulatory networks [3]). Even when the biological system does not have a network structure, a network model can be leveraged by treating an empirical distance or similarity matrix (e.g., covariance matrix for gene expression data or physical distance matrix for spatial data) as a network adjacency matrix. In this short paper, we explore techniques for analyzing biological networks using the Krylov subspace computed on the network adjacency matrix. In particular, we focus on the scenario where a biological state or perturbation of interest can represented by a vector of node-specific weights and this vector, rather than a random vector, is used to initialize the Krylov subspace computation. The order *m* Krylov subspace [4, 5] for a square diagonalizable *p × p* matrix **X** and length *p* vector **v** corresponds to the sequence of *m* vectors generated by the product of **v** with powers of **X** ranging from 0 to *m* − 1, i.e. {**v, Xv, X**^**2**^**v**, …, **X**^**m**−**1**^**v**}. This sequence of vectors can be computed via the method of power iteration with per iteration normalization [6] and used to form the columns of a *p × m* Krylov subspace matrix **K**:

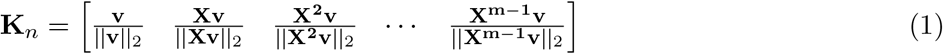

The Krylov subspace is leveraged in a diverse range of numerical linear algebra techniques. In particular, the columns of **K** usually converge to the eigenvector of **X** associated with the largest eigenvalue as long as **v** is not orthogonal to this principal eigenvector. The columns of **K** are also linearly independent assuming *m* ≤ *r* + 1 where *r* is the rank of **X**. These properties enable methods such as Arnoldi iteration [7], which computes the eigenvectors and eigenvalues of **X** using the Krylov subspace with incremental orthogonalization, Lanczos iteration [8], the conjugate gradient method [9] and the minimal residual method [10].

In the network analysis space, the Krylov subspace of **A** is most commonly generated by executing power iteration until convergence with a random initial vector to generate eigenvector centrality values. For strongly connected networks, **A** is irreducible and the Perron-Frobenius theorem [11] ensures there is a unique largest real eigenvalue and corresponding eigenvector with strictly positive elements (the eigenvector centrality values). In this scenario, only the final column of **K** is of interest and the rest are discarded. One network analysis method that does leverage the full Krylov subspace of **A** is the flow profiles method of Cooper and Barahona [12], which computes node similarity values for directed networks by calculating a matrix **X** that combines the columns of the scaled Krylov subspace matrices for both **A** and **A**^*T*^ using a uniform initial vector set to **1** (vector of all ones). Note that for directed networks, power iteration performed on **A** and **A**^*T*^ will converge to different final vectors with distinct intermediate results, which implies that there are two Krylov trajectories for each node of a directed network. While the discussion in most of the manuscript focuses on analysis of symmetric adjacency matrices, the fact that the **A** and **A**^*T*^ matrices associated with directed networks have distinct Krylov subspaces is leveraged by the flow profiles method and in our analysis of the C. Elegans neural network. The network path intepretation of the Krylov subspace of **A** is also related to the node similarity measure proposed by Leicht, Holme and Newman (LHN) [13], which uses a matrix defined in terms of the powers of **A**. In particular, the *r*^*th*^ column of **K** takes the form **K**[, *r*] = **A**^*r*^**v**. Since the elements of **A**^*r*^ represent the weighted count of paths of length *r* between each node pair, this implies that element *i* of the column **K**[, *r*] is proportional to the weighted count of all paths of length *r* between node *i* and all other nodes in the network.

In this paper, we focus on the Krylov subspace of a biological network adjacency matrix **A** as computed by power iteration using a non-random and potentially non-uniform initial vector **v** that encodes a biological state or perturbation. This investigation extends our earlier work exploring the general properties of the Krylov subspace of **A** using uniform initial vectors [14]. Specifically, this paper introduces two functions of the Krylov trajectories that capture gradiant information (Krylov velocity vectors) and node importance (the *δ* statistic) and explores the utility of these statistics for community detection and perturbation analysis using the C. Elegans neural network.

## 2 Methods

### 2.1 Krylov trajectories, velocity vectors and *δ* statistics

We are interested in the order *m* Krylov subspace matrix **K** associated with a network adjacency matrix **A** (and **A**^*T*^ for directed networks) as computed via *m* − 1 power iterations on **A** using a non-random initial vector **v**. We refer to the rows of **K** as Krylov trajectories and represent the trajectory for node *i* by **t**_*i*_:

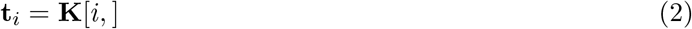

We are also interested in an approximation of the gradient of the Krylov trajectories that we refer to as Krylov velocity vectors and represent by 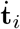. These velocity vectors are formed by replacing the elements in each Krylov trajectory vector with the difference from the prior element (or 0 for the initial element):

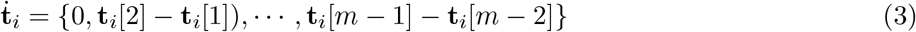

The veolocity vectors represent an embedding of each node that explicity encodes the ordering of the columns of the Krylov subspace matrix.

Krylov trajectories can also be used to capture node importance. Since the last value of each trajectory captures eigenvector centrality, we are specifically interested in functions of the entire trajectory that capture aspects of node importance distinct from eigenvector centrality. Although many possible functions exist, we believe that functions which quantify the level of oscillations in the trajectory may have particular utility. One simple way to quantify this is to subtract the absolute difference between the starting and ending values of a given trajectory from the sum of the absolute values of 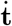 (i.e., the absolute values of all sequential differences). We refer this trajectory statistic for node *i* as *δ*_*i*_:

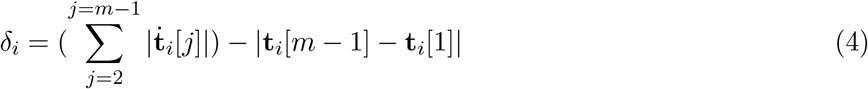

The *δ* statistic captures the number and magnitude of oscillations in the trajectory (for monotonic trajectories, it is 0):

### 2.2 Node similarity and clustering using Krylov trajectories

One of the key applications of the Krylov trajectories and velocity vectors is the computation of node similarity values, which can be leveraged for clustering/community detection. Importantly, node clustering using trajectory-based similarity values yields very distinct results from standard approaches for network community detection such as Louvain clustering [15] or clustering using Leicht, Holme and Newman (LHN) similarity values [13]. To illustrate this phenomenon, we performed clustering using three different methods: 1) Louvain clustering as implemented by the *igraph* R package [16] function *cluster louvain()*, 2) hierarchical agglomerative clustering (the dendrogram is cut to match Louvain clustering results) using a distance of one minus the Pearson correlation between Krylov trajectories or velocity vectors, and 3) hierarchical agglomerative clustering using a distance of one minus the LHN regular equivalence between nodes (as computed using the *linkprediction* R package function *proxfun(method=“lhn global”)* [17]). The dendrogram is again cut to match Louvain clustering.

### 2.3 Perturbation analysis using Krylov trajectories

To explore the impact of a specific perturbation (or other biological condition), a non-uniform intial vector with node-specific weights can be used. Although power iteration will converge to the same value irrespective of the initial **v**, the Krylov trajectories (and 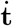 and *δ* statistics) will capture the impact of the perturbation on the network.

## 3 Results

To explore the practical utility of Krylov trajectories, velocity vectors and the *δ* statistic, we analyzed the C. Elegans neural network compiled by Chen et al. [18] and downloaded from the WormWeb project (neural connections from http://wormweb.org/data/NeuronConnect.txt and neuron types from http://wormweb.org/data/name_neurons.txt). Simulations studies were also performed but are not included given length constraints. Although more comprehensive C. Elegans networks have been compiled that include connections to non-neuronal cells [19], we choose to focus on the less complex network from Chen et al. capturing just nonpharyngeal neurons. After removing neurons lacking type information, this resulted in a directed and unweighted network with nodes representing 275 neurons and 3,606 edges representing neural connections. Neuron types were classified as sensory, interneuron or motor neuron with some neurons having multiple types. Neuron connections were specified as either electrical junctions (treated as bidirectional edges) or chemical synapses (treated as directed edges). This network is visualized in the top left panel of Fig 1.

**Figure 1.**
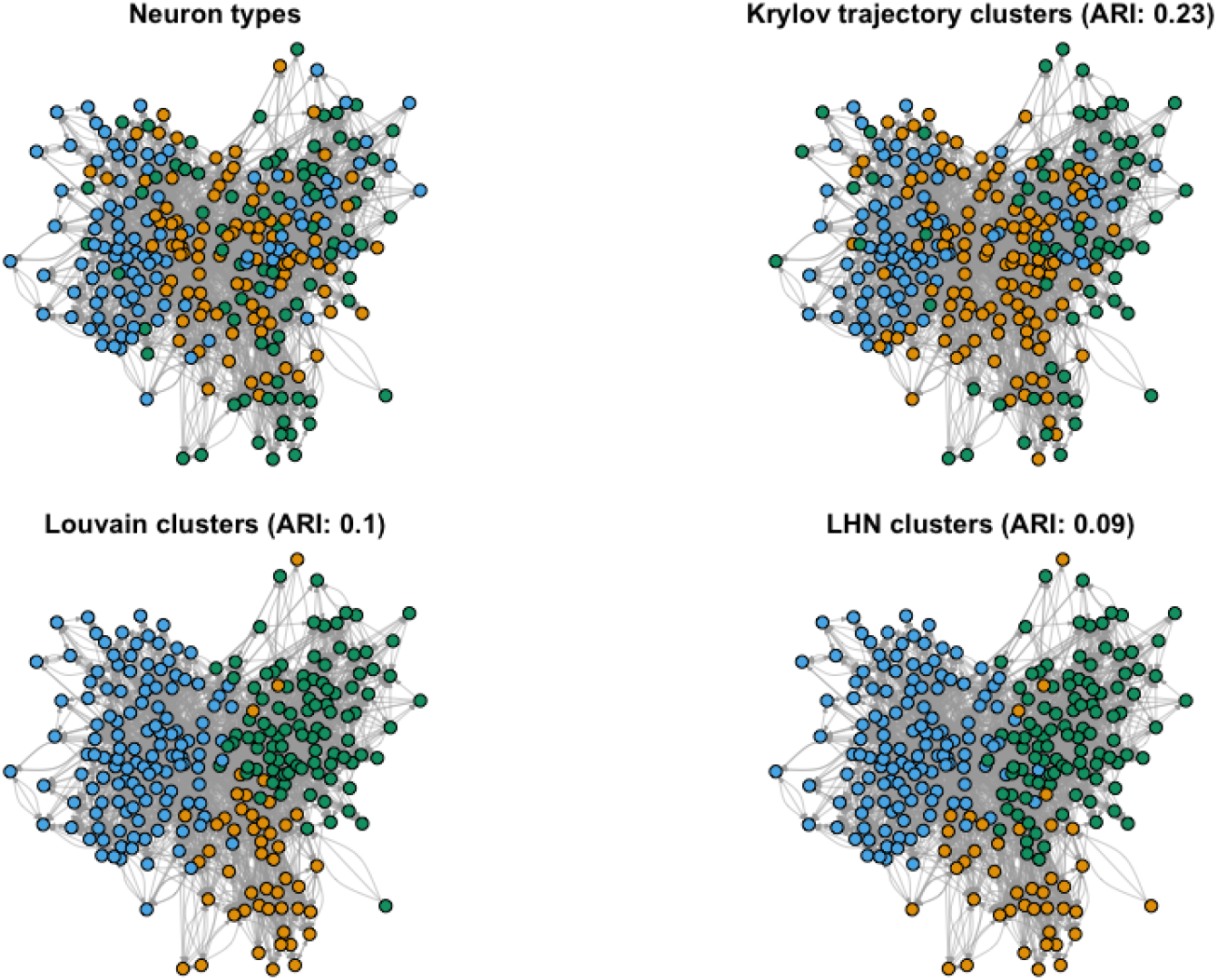
Visualization of the C. Elegans neural network. The top left panel visualizes neuron type (green: sensory, blue: motor, orange: interneuron). For neurons assigned multiple types, the node is colored according to the first listed type. The other panels visualize the results of clustering according to Krylov trajectory vectors, Louvain clustering or LHN similarity values with corresponding Adjusted Rand Index (ARI) values in parentheses.

### 3.1 Identification of neuron type groups

As an initial analysis, we performed clustering on this C. Elegans neural network using Krylov trajectories, Louvain clustering, and LHN node similarity values. Both Louvain and LHN-based clustering were performed on a undirected version of the network following the approach outlined in Section 2.2. For Krylov trajectory clustering, trajectories were created using both **A** and **A**^*T*^ (i.e., the adjacency matrix and its transpose) and these two trajectory vectors were combined to form a single vector for each node that was then employed to compute the correlation-based node similarity values used for hierarchical agglomerative clustering. All methods were executed to yield three clusters matching the number of distinct neuron types with the results and the associated Adjusted Rand Index (ARI) values shown in Fig 1. Although none of the methods generated high ARI values, the use of Krylov trajectory vectors performed noticably better than either Louvain or the LHN-based method. The poor performance of community detection on this network may simply indicate that there is only a limited association between neuron type and network structure in C. Elegans, however, it is possible that classification performance is impacted by various approximations made when creating the network. These approximations include ambigious neuron type labels (multiple neurons are assigned two types), the fact that neural connection weights were not included, the incorporation of both electrical junctions and chemical synapses in a the same network, and fact that the network does not include connections to non-neural cells.

### 3.2 Response to sensory neuron stimulation

To explore the ability of Krylov trajectory vectors to characterize network perturbations, we computed the *δ* statistic for each node on the non-perturbed network and with a perturbation applied to the left and right-side ADE sensory neurons (ADEL and ADER). Similar to the use of Krylov trajectory vectors for clustering, the *δ* statistic was computed on both **A** and **A**^*T*^ and the sum of these two values was used as the final *δ* statistic. The perturbation was specifically modeled using an initial vector **v** where the values for ADEL and ADER are 10 times larger than the values for other nodes. The top panels in Fig 2 visualize the *δ* values for both the unperturbed and perturbed cases with *δ* differences shown in the bottom panels. The neurons with the 10 largest *δ* values for both cases are listed in Table 1. For the unperturbed network, motor and interneurons in general have larger *δ* values than sensory neurons, which is consistent with an expectation that these neurons are likely to be more sensitive to neural perturbations. The targeted perturbation of the ADEL/ADER sensory neurons results in a noticable increase in *δ* values for a small group of interneurons (e.g., RIH and RIGL/RIGR), which presumably represent the interneurons that are most sensitive to ADE-mediated sensory inputs. Interesting, the perturbed *δ* values show a right/left asymmetry with larger values for the right-side neurons, which is consistent with the known left/right asymmetry of neural responses in C. Elegans [20].

**Table 1:** Neurons with the 10 largest *δ* values in both the unperturbed and perturbed networks.

| Neuron | Unperturbed $\delta$ | Neuron | Perturbed $\delta$ |
| --- | --- | --- | --- |
| AVAR | 0.292 | AVAR | 0.427 |
| AVAL | 0.291 | AVAL | 0.328 |
| AS7 | 0.186 | AVKR | 0.248 |
| AS9 | 0.152 | RIH | 0.236 |
| AVBR | 0.150 | RIGR | 0.186 |
| AIBR | 0.146 | RIAL | 0.186 |
| DB5 | 0.141 | AVKL | 0.173 |
| DB6 | 0.137 | AS7 | 0.168 |
| AVER | 0.137 | RIGL | 0.167 |
| AS8 | 0.135 | AS8 | 0.151 |

**Figure 2.**
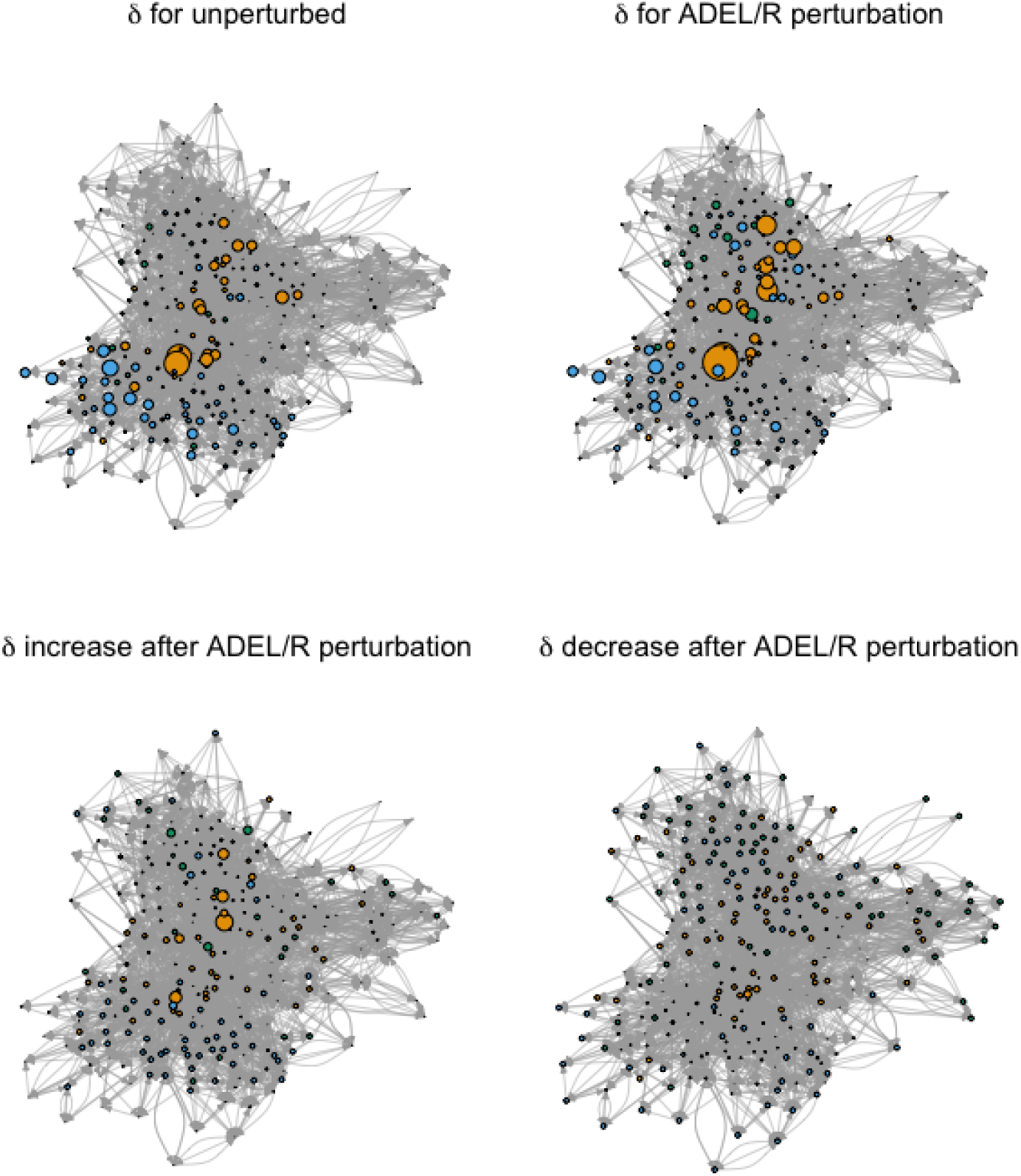
Visualization of the *δ* statistic for the unperturbed C. Elegans neural network and the network with a perturbation to the ADEL and ADER sensory neurons.

## Acknowledgments

This work was funded by National Institutes of Health grants R35GM146586, P20GM130454 and P30CA023108. We would like to acknowledge the supportive environment at the Geisel School of Medicine at Dartmouth where this research was performed.

